# Grape branch substrates promote adaptive glycerophospholipid metabolism and lipid-synthesis gene expression in *Stropharia rugosoannulata* under moderate nitrogen limitation

**DOI:** 10.64898/2026.11.15.763823

**Authors:** Kexin Xie, Meng Shen, Hongkai Wang, Qunli Jin, Weiming Cai, Fucheng Lin, Xiaohong Liu

## Abstract

This study investigated the use of lignified grape plant materials (branches, stems, and roots) as cultivation substrates for *Stropharia rugosoannulata* and their effects on fruiting-body lipid metabolism. Metabolomic profiling identified glycerophospholipid metabolism as the primary pathway showing differential responses among substrates. While overall lipid composition remained largely unchanged, intermediate Lyso-GPL metabolites (LPA, LPC, LPI, and LPS) were more active under grape-branch substrates, indicating enhanced membrane-lipid metabolic remodeling. Simulation experiments with varying C/N ratios demonstrated that moderate nitrogen limitation (C/N = 28:1) significantly upregulated key lipid-synthesis genes (ACS1, ACS2, ACC, and FAS), with ACS1 showing the strongest response, and also activated the transcription factor GCN4. Under severe nitrogen limitation (C/N = 64:1), ACS1 upregulation persisted but GCN4 activation did not, suggesting a transition toward metabolic suppression. Further TOR-specific inhibition experiments showed that continuous TOR inactivation (24 h) sustained high GCN4 expression while downregulating ACS1 and ACS2, with ACC remaining relatively stable. Genetic manipulation confirmed that GCN4 positively regulates tolerance to oxidative and cell-wall stress, whereas its continuous overexpression sensitized mycelia to amino-acid starvation (3-AT). These findings indicate that moderate nitrogen restriction, characteristic of grape-branch substrates, preferentially engages adaptive lipid-metabolic regulatory networks involving GCN4 that may improve mushroom quality.

## Introduction

*Stropharia rugosoannulata* is an edible mushroom of increasing commercial interest that can be cultivated on a wide range of lignocellulosic agricultural wastes [1-3]. In grape-producing regions, pruned branches, stems, and roots represent abundant, underutilized lignified materials that could serve as low-cost cultivation substrates while contributing to circular agriculture. However, the effects of these grape-derived substrates on the metabolic quality of the resulting fruiting bodies remain poorly understood.

Lipid composition is an important determinant of mushroom flavor, texture, and nutritional value [4-6]. Previous studies have shown that substrate carbon-to-nitrogen (C/N) ratio can influence fungal metabolism[7,8], yet the specific impact of grape-derived lignified materials on lipid metabolic pathways in *S. rugosoannulata* has not been systematically examined. In particular, it is unclear whether differences among substrates primarily alter terminal storage lipids or act through intermediate metabolic steps, and whether moderate nitrogen limitation, which is common in highly lignified materials, triggers adaptive transcriptional responses.

In this study, we used grape branches, stems, and roots as cultivation substrates and applied metabolomic and gene-expression analyses to investigate how these materials affect lipid accumulation and glycerophospholipid metabolism in the fruiting bodies of *S. rugosoannulata*. Special attention was paid to Lyso-GPL intermediates and the potential regulatory role of the transcription factor GCN4 under different degrees of nitrogen limitation. To further dissect the signaling mechanisms underlying nitrogen-limitation responses, we examined the effects of continuous TOR inhibition on GCN4 and lipid-metabolism genes, and functionally validated the role of GCN4 in amino-acid starvation and environmental stress tolerance through overexpression and RNAi approaches.

## Materials and Methods

### Strains and culture conditions

The monokaryotic *S. rugosoannulata* strain and the dikaryotic strain were previously generated and characterized in our study. All *S. rugosoannulata* strains were routinely cultured on complete yeast medium (CYM), potato dextrose agar (PDA), or minimal medium (MM) as indicated.

### Nitrogen gradient / nitrogen-limitation treatments

Carbon-to-nitrogen (C/N) ratio gradient induction: Gradient media ranging from nitrogen-replete to nitrogen-limited conditions were prepared with C/N ratios of 8:1 (high-nitrogen control with sufficient nitrogen supply), 28:1 (moderate nitrogen limitation), and 64:1 (severe nitrogen limitation). Dikaryotic mycelia were cultured under these conditions, and samples were collected at different growth stages for phenotypic observation and gene expression analysis.

Nitrogen reduction and acute nitrogen starvation treatments: High-nitrogen medium (HN) was used as the control. Low-nitrogen (LN) and complete nitrogen-free / acute nitrogen starvation (−N) conditions were established. Dikaryotic mycelia were subjected to these nutrient-limitation treatments to compare metabolic reprogramming under chronic nitrogen limitation versus acute nitrogen starvation.

### Total RNA extraction and reverse transcription-quantitative PCR (RT-qPCR)

Dikaryotic mycelia from each treatment were collected, flash-frozen in liquid nitrogen, and ground to a fine powder. Total RNA was extracted, genomic DNA was removed by DNase treatment, and cDNA was synthesized by reverse transcription. Specific primers were designed for the amino-acid starvation-responsive transcription factor gene GCN4 and for core lipid-metabolism genes (FAS, ACC, ACS1, ACS2, ACL1, and DGAT2). RT-qPCR was performed, and relative expression levels were calculated using the 2−ΔΔCt method, with the high-nitrogen group (HN or C/N 8:1) serving as the calibrator. Biological and technical replicates were included for each sample. Primer information is provided in Table 1. All gene sequences are provided in Yu et al. (2026) [9].

**Table 1.**
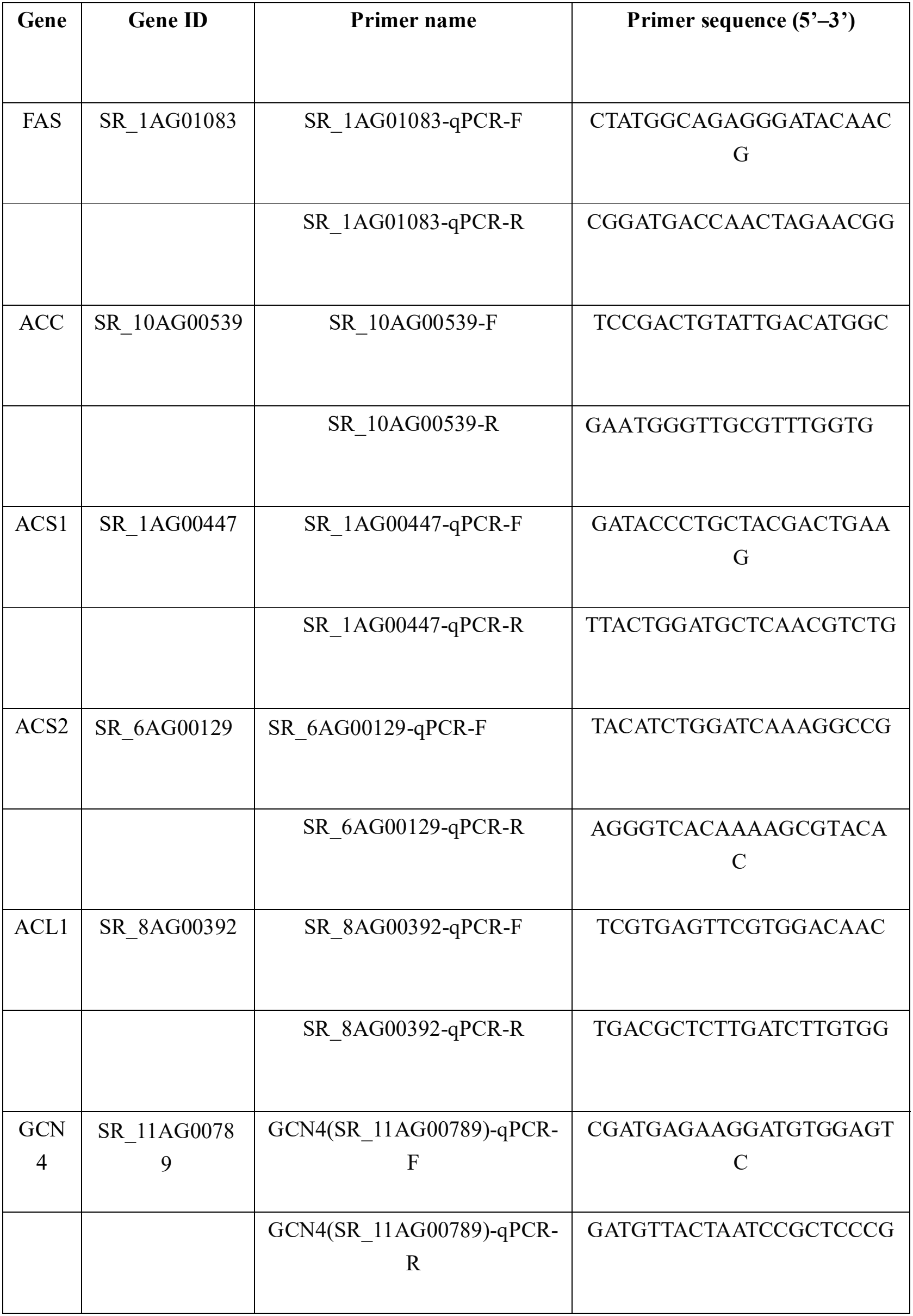
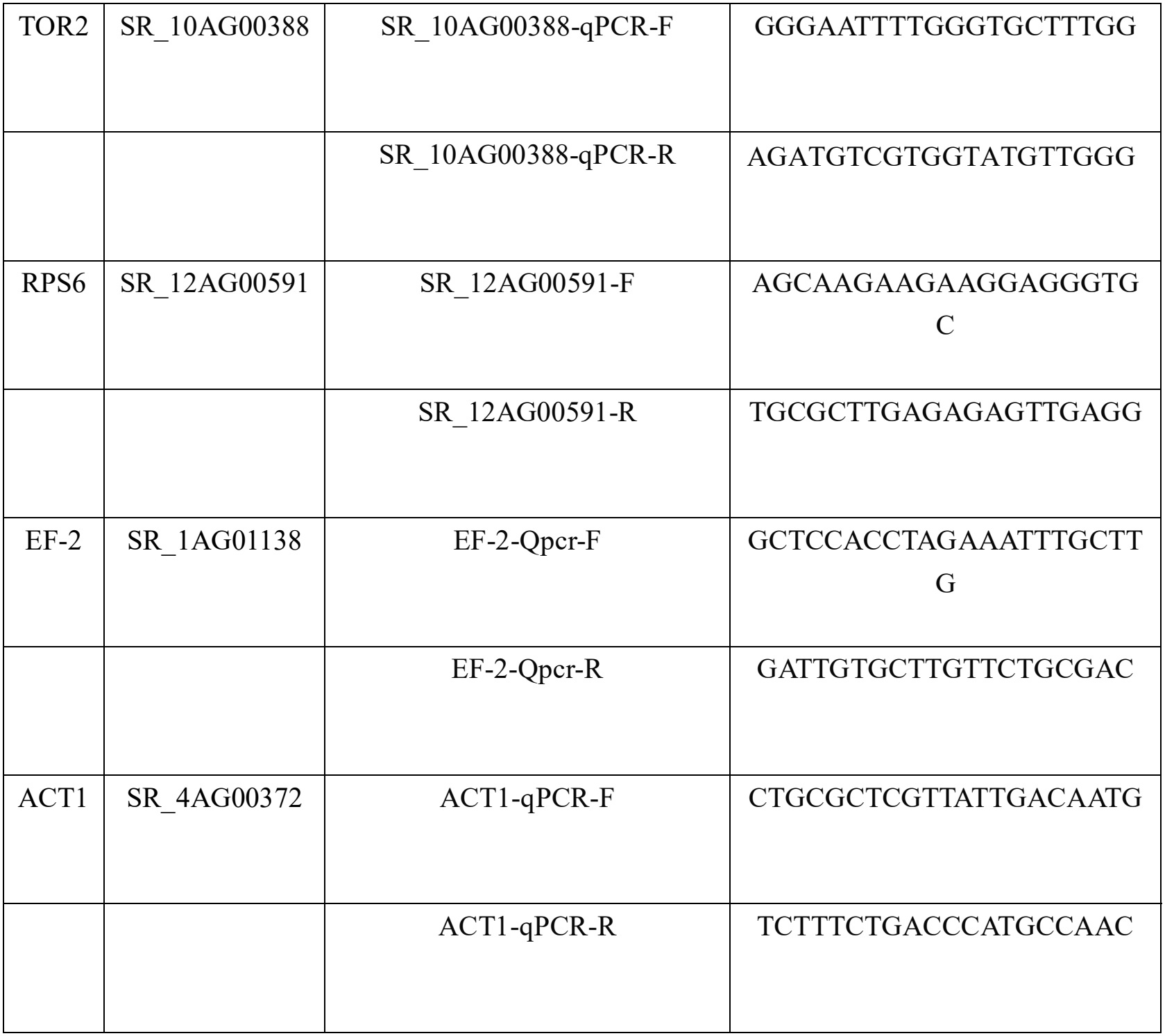
Gene ID and Primers for this study.

### Construction of GCN4 overexpression and RNAi strains and multi-stress tolerance assays

GCN4 overexpression (OE) transformants (e.g., g-6) and RNAi knockdown transformants (e.g., g-2-14 and g-2-18) were generated. Wild-type (WT) strain was used as the control. Transformants were inoculated onto solid media supplemented with different exogenous stress agents, and colony diameters were measured to calculate relative growth rates.

Amino-acid starvation stress: 3-Amino-1,2,4-triazole (3-AT), a competitive inhibitor of histidine biosynthesis, was added to mimic intracellular amino-acid starvation.

Oxidative stress: H□O□was added to assess reactive oxygen species (ROS) scavenging capacity and oxidative stress tolerance.

Cell-wall integrity stress: Congo Red was added to evaluate cell-wall stress resistance and repair capacity.

## Results

### Effects of grape-derived lignified materials (branches, stems, and roots) as cultivation substrates on the quality of *S. rugosoannulata*

In this phase of the study, lignified materials of different origins from grape plants (branches, stems, and roots) were used as cultivation substrates, and their effects on the lipid accumulation characteristics of the fruiting bodies of *Stropharia rugosoannulata* were systematically analyzed. Metabolomic analysis revealed that the most pronounced differences in the response of *S. rugosoannulata* to the various substrates were concentrated in the glycerophospholipid metabolism pathway (Fig. 1). Although the overall lipid compositional proportions did not change substantially, clear differential accumulation was observed at the metabolic level, particularly among glycerophospholipid-related intermediate metabolites, whereas the terminal storage lipids remained relatively stable. These findings indicate that substrate differences primarily regulate intermediate steps of lipid metabolism rather than directly altering the final lipid storage forms (Fig. 2).

**Fig. 1.**
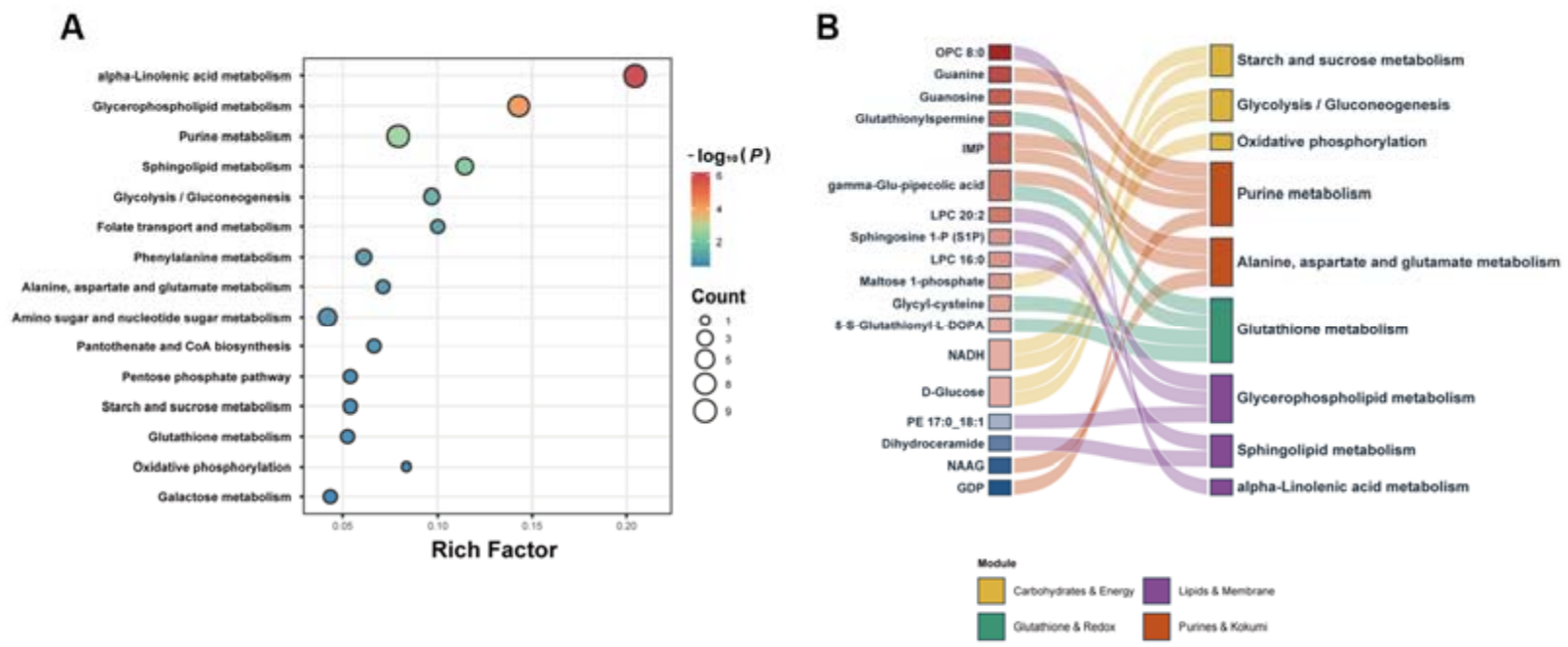
(A) KEGG enrichment bubble plot of differential metabolites between the Branch and Wood groups. The x-axis represents the enrichment factor (Rich Factor), bubble size indicates the number of differential metabolites enriched in the pathway (Count), and bubble color represents the enrichment significance level (−log□□P). (B) Sankey diagram showing the topological associations between 17 representative differential metabolites (left nodes, colored from blue to red according to the magnitude of Branch/Wood log□FC changes) and 9 core KEGG metabolic pathways (right nodes, colored by four major functional modules).

**Fig. 2.**
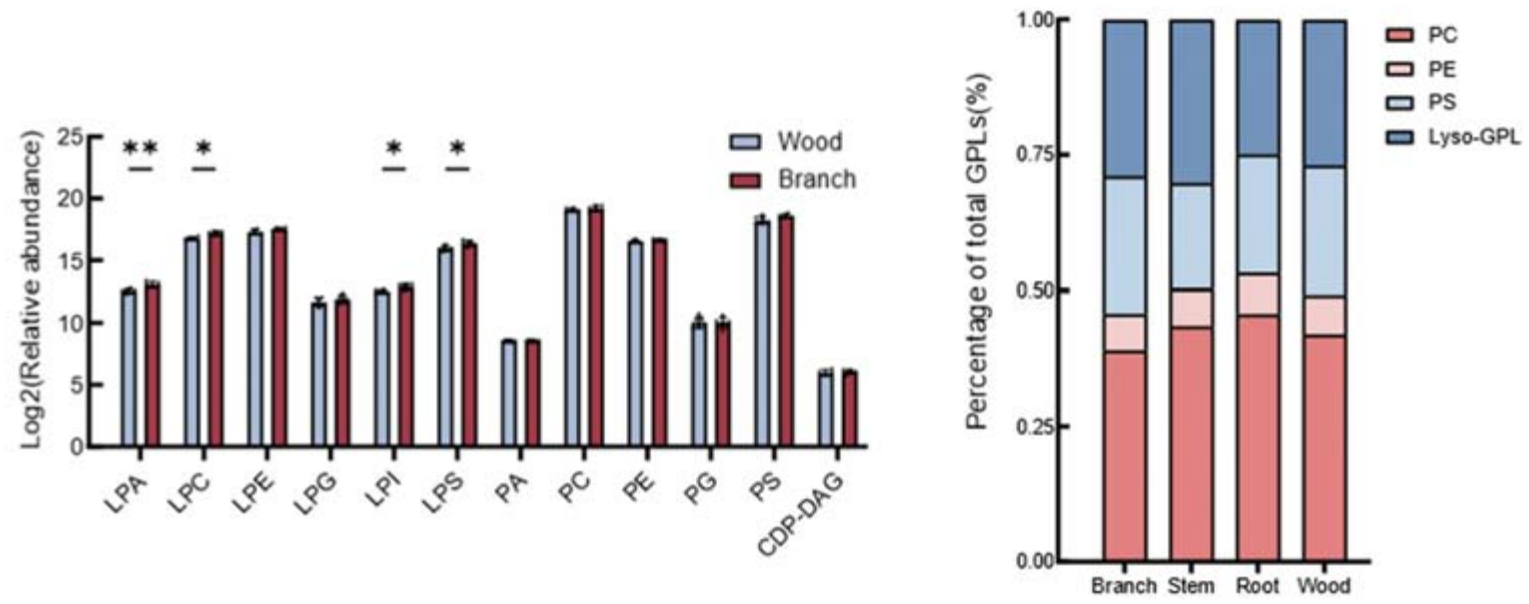
Metabolic activity of Lyso-GPL intermediates and overall lipid composition under grape-branch-related substrates. In substrates derived from grape branches, Lyso-GPL-type intermediates (LPA, LPC, LPI, and LPS) exhibited higher metabolic activity, whereas the overall proportional structure of lipids showed no significant changes.

Further analysis showed that, under substrate conditions represented by grape branches, Lyso-GPL-type intermediates (including LPA, LPC, LPI, and LPS) exhibited higher metabolic activity. This suggests that such substrates more strongly drive membrane-lipid-related metabolic remodeling. When these observations were integrated with simulation experiments using different carbon-to-nitrogen (C/N) ratios, multiple key genes involved in lipid synthesis (including *ACS1, ACS2, ACC*, and *FAS*) were significantly upregulated under moderate nitrogen limitation (C/N = 28:1, corresponding to the actual grape-branch substrate). Among these, the upregulation of *ACS1* was particularly pronounced (Fig. 3). Concurrently, the transcription factor GCN4 was significantly activated under this condition, suggesting that it may mediate adaptive metabolic regulation in response to nitrogen limitation.

**Fig. 3.**
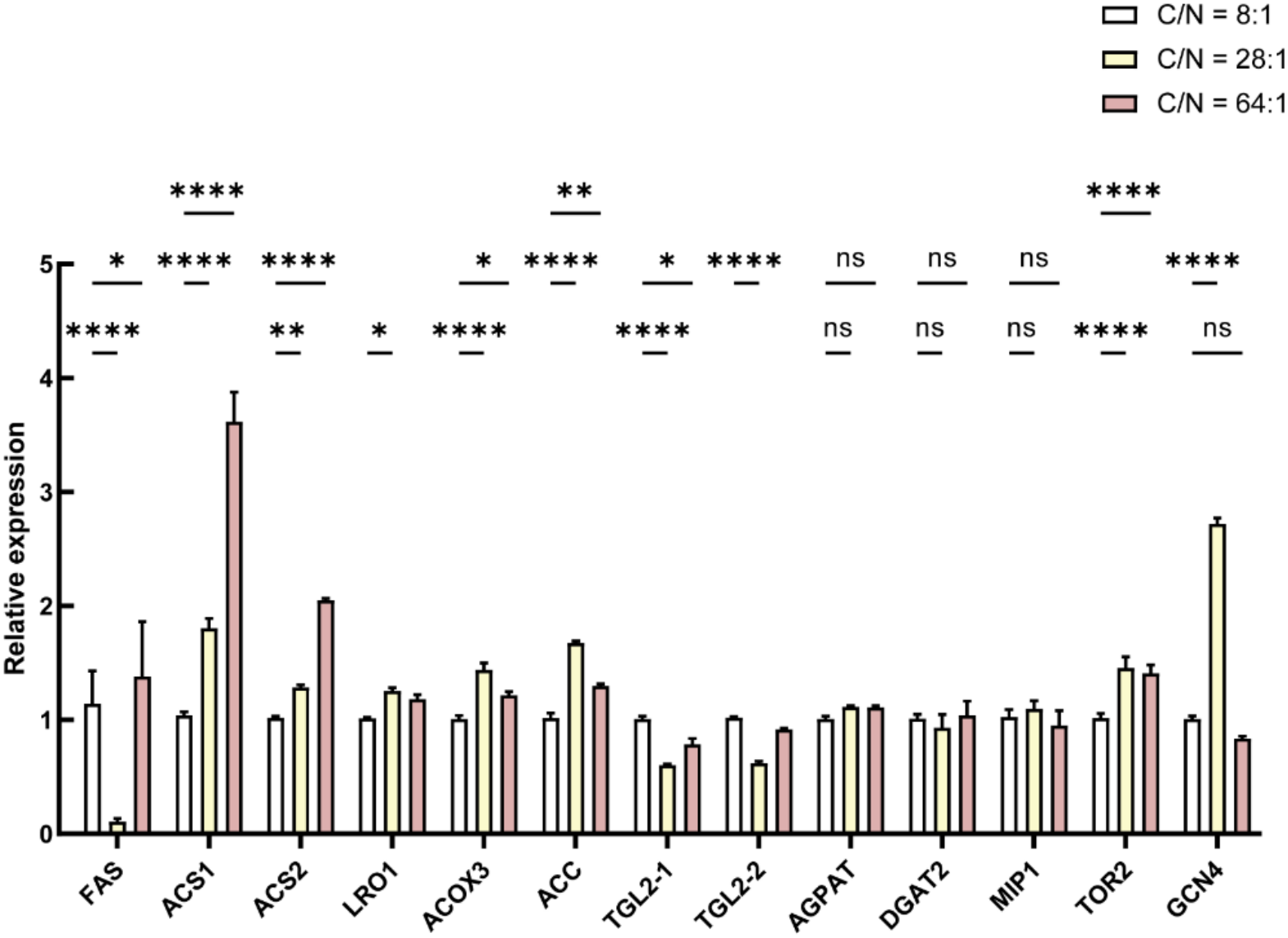
Relative expression levels of lipid metabolism-related genes (GCN4, FAS, ACS1/ACS2, ACC, etc.) under the different carbon-to-nitrogen (C/N) ratio gradients (8:1, 28:1, 64:1) and treatment conditions.

In contrast, under strong nitrogen limitation (C/N = 64:1, simulating highly lignified or late-stage substrates), *ACS1* remained upregulated, but GCN4 expression was not further enhanced. This pattern implies that the system may have entered a state of metabolic inhibition or resource depletion. Under low C/N conditions (8:1, laboratory control), no comparable regulatory pattern was observed. Collectively, these results indicate that a moderate degree of nitrogen limitation is more favorable for activating the adaptive regulatory network associated with lipid metabolism. The specific functional role of GCN4 in this process is currently being verified through further experiments.

### Dynamic balance of GCN4, ACS1/ACS2, and ACC under continuous TOR inhibition

To clarify the upstream and downstream relationships of the TOR (Target of Rapamycin) pathway in nitrogen-limitation-induced lipid metabolic reprogramming, mycelia were subjected to TOR-specific inhibition treatment, and samples were collected at time points including 24 h of continuous inhibition. The dynamic expression responses of the core stress transcription factor (GCN4), the ribosomal protein gene (RPS6), and downstream fatty acid synthesis- and activation-related genes (ACC, ACS1, ACS2, FAS, etc.) were then examined.

Gene expression profiles after 24 h of continuous TOR inhibition were analyzed, with TOR2 representing the TOR pathway itself, GCN4 and RPS6 reflecting stress and translational status, ACS1/ACS2 assessing fatty acid metabolic flux, and ACC serving as a benchmark indicator of fatty acid synthesis capacity. GCN4 showed sustained activation. TOR inhibition rapidly and significantly induced high expression of GCN4, indicating that TOR inactivation directly triggers nutrient stress and amino acid starvation responses. Under continuous 24 h inhibition, GCN4 maintained high-level expression.

In contrast, ACS1 and ACS2 showed downregulated responses. Under 24 h continuous TOR inhibition, the expression levels of ACS1 and ACS2 were markedly downregulated, suggesting that under prolonged extreme stress lacking growth signals, the overall metabolic flux of free fatty acid activation and downstream lipid turnover decreased as cells reduced energy-intensive metabolic activities.

ACC maintained steady-state levels. During continuous TOR inhibition, the transcript level of ACC remained essentially stable (ACC ≈ unchanged), indicating that even under growth-suppressed conditions, cells maintained the basal rate-limiting enzyme capacity for de novo fatty acid synthesis at a baseline level (Fig. 4) to ensure a smooth transition from a growth-oriented state to metabolic reprogramming and energy storage states.

**Fig. 4.**
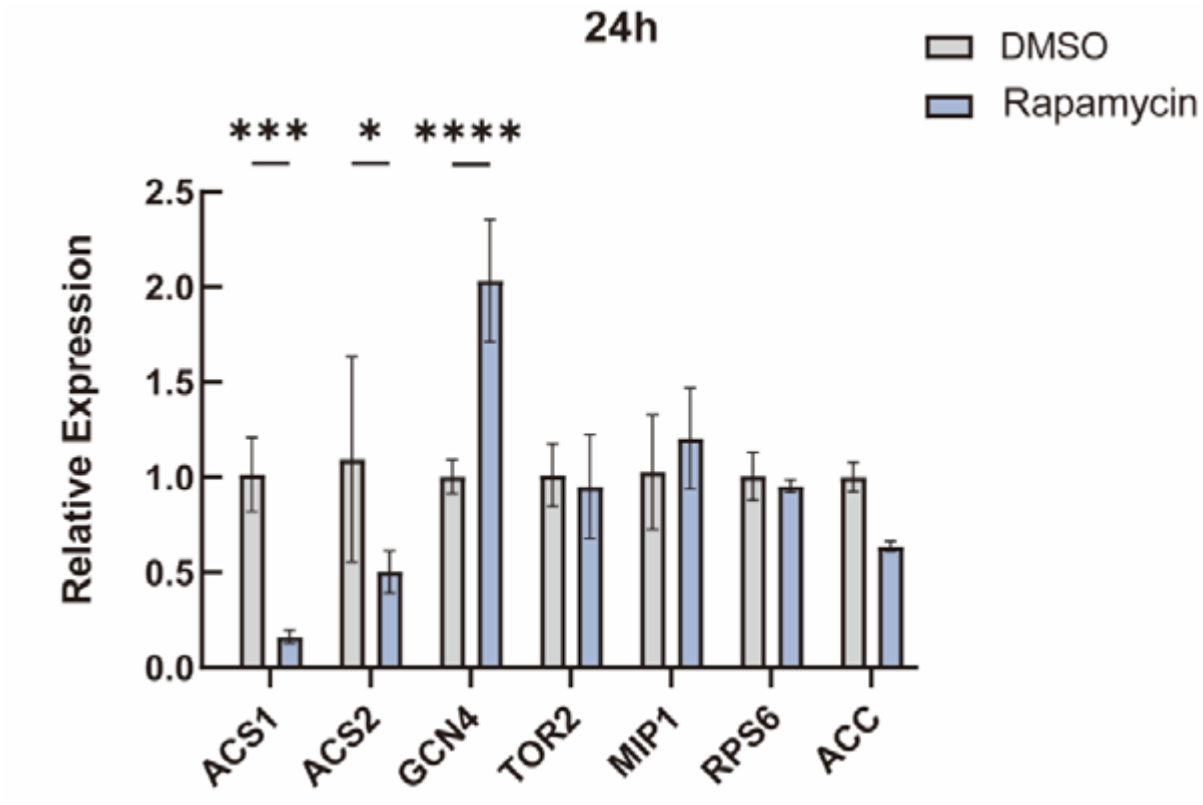
Relative expression changes of TOR2, GCN4, RPS6, and lipid metabolism genes ACS1, ACS2, and ACC under continuous TOR inhibition (24 h).

### Functional verification of GCN4 genetic manipulation (overexpression/interference) in regulating amino acid starvation and environmental stress

To further verify the biological function of GCN4 in nutrient starvation and stress adaptation, GCN4 overexpression transformants and RNAi interference transformants were successfully constructed (Fig. 5), and the growth phenotypes of each strain under different stress conditions were determined. Phenotypic functional verification of the wild-type (WT), GCN4 overexpression strains (OE), and interference strains (RNAi) further clarified the central role of GCN4 in nutrient starvation and stress adaptation.

**Fig. 5.**
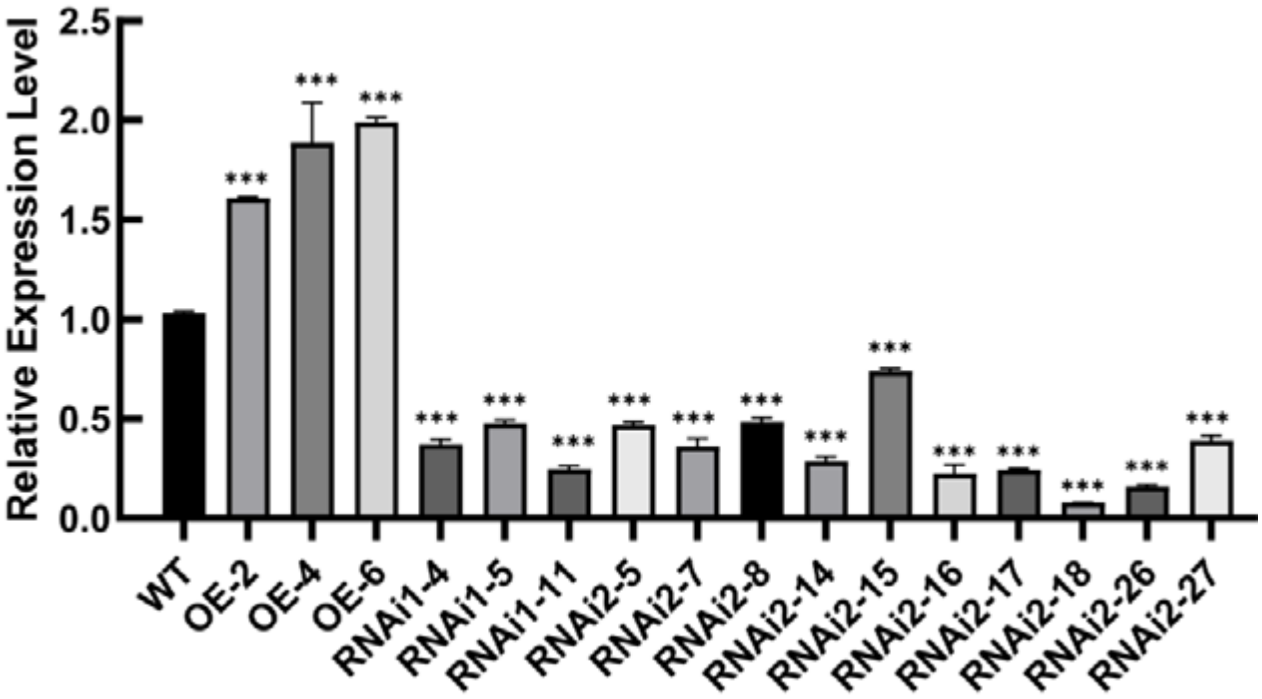
Construction and screening verification of GCN4 overexpression (OE) and RNAi interference transformant strains.

Differential sensitivity to 3-AT (amino acid starvation stress) was observed. 3-AT inhibits histidine synthesis and induces intracellular amino acid starvation, producing growth inhibition in all tested strains. However, the degree of response differed significantly among strains with different GCN4 expression levels: colony diameter of GCN4 overexpression strains was markedly reduced under 3-AT stress, with overexpression strain g-6 being the most sensitive; conversely, some GCN4 interference strains (such as g-2-14 and g-2-18) showed higher relative growth rates under 3-AT and were less inhibited (Fig. 6). This indicates that exogenous continuous overexpression of GCN4, when superimposed with the strong starvation signal of 3-AT, may trigger excessive metabolic reprogramming arrest and growth braking effects.

**Fig. 6.**
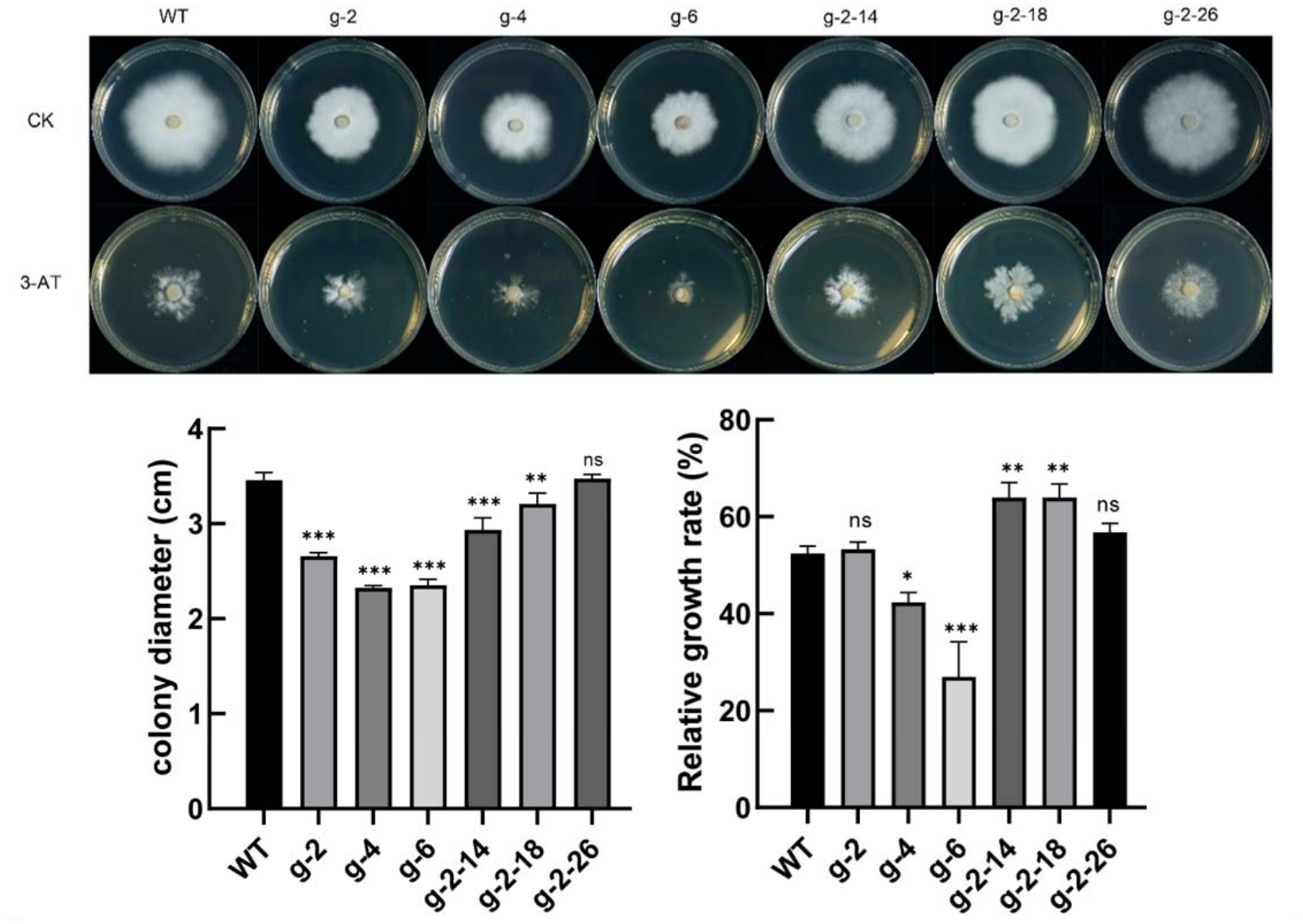
Colony growth phenotypes and relative growth rate comparison of WT, GCN4 overexpression strains, and interference strains under 3-AT (amino acid starvation) stress.

Positive regulation of oxidative stress and cell wall stress was also confirmed. In plate assays with H□O□(oxidative stress) and Congo Red (cell wall stress), GCN4 overexpression significantly enhanced the tolerance of strains to H□O□and Congo Red, whereas GCN4 interference strains showed markedly reduced tolerance. This confirms that GCN4 positively regulates the antioxidant stress and cell wall integrity maintenance pathways of mycelia (Fig. 7).

**Fig. 7.**
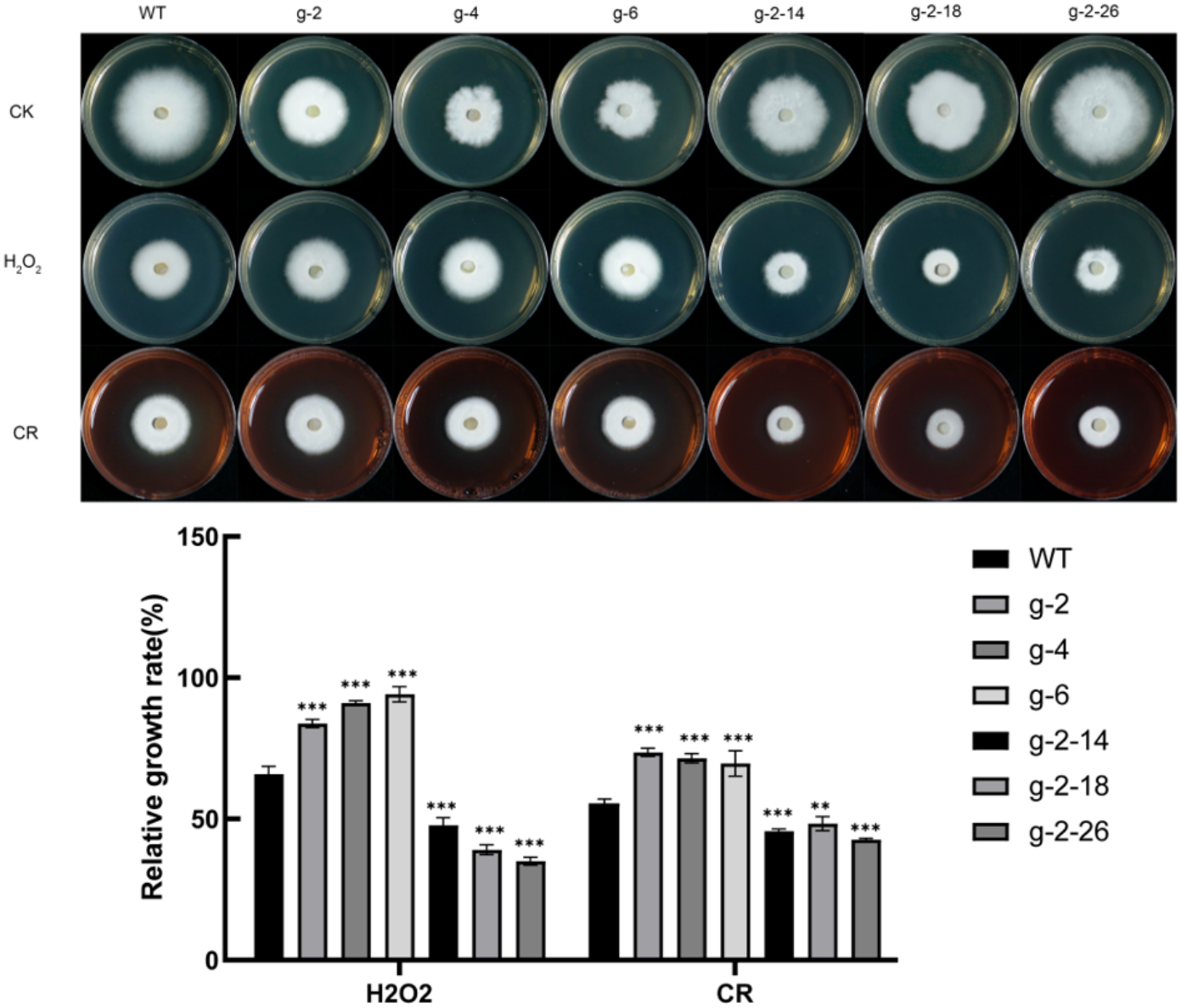
Tolerance phenotype analysis of WT, GCN4 overexpression strains, and interference strains under H□O□ (oxidative stress) and Congo Red (cell wall stress) conditions.

## Conclusions

This study demonstrates that lignified grape-derived materials (especially branches) can serve as effective cultivation substrates for *S. rugosoannulata* by modulating lipid metabolism primarily at the intermediate rather than terminal storage level. Metabolomic analysis showed that glycerophospholipid metabolism, particularly Lyso-GPL intermediates (LPA, LPC, LPI, and LPS), was the core pathway responding to substrate differences, while overall lipid compositional proportions remained relatively stable.

Moderate nitrogen limitation (C/N = 28:1), characteristic of grape-branch substrates, preferentially activated adaptive lipid-metabolic regulation, including significant upregulation of key lipid-synthesis genes (*ACS1, ACS2, ACC*, and *FAS*) and the transcription factor GCN4. In contrast, severe nitrogen limitation (C/N = 64:1) maintained *ACS1* upregulation but failed to sustain GCN4 activation, suggesting a shift toward metabolic suppression. Continuous TOR inhibition further revealed that TOR inactivation directly triggers sustained GCN4 expression while downregulating ACS1 and ACS2, with ACC remaining relatively stable, indicating a coordinated reprogramming of lipid metabolism under growth-inhibitory conditions.

Functional verification through GCN4 overexpression and RNAi confirmed that GCN4 positively regulates mycelial tolerance to oxidative stress and cell-wall stress. However, continuous GCN4 overexpression sensitized strains to amino-acid starvation induced by 3-AT, indicating a context-dependent role of GCN4 in balancing stress adaptation and growth.

Collectively, these findings indicate that moderate nitrogen restriction associated with grape-branch substrates engages GCN4-mediated adaptive regulatory networks that remodel membrane lipid metabolism, potentially contributing to improved fruiting-body quality. The results provide a molecular basis for the rational utilization of grape pruning residues in *S. rugosoannulata* cultivation.

